# Increased baseline RASGRP1 signals Enhance Stem Cell Fitness during Native Hematopoiesis

**DOI:** 10.1101/2020.06.24.169631

**Authors:** Laila Karra, Damia Romero-Moya, Olga Ksionda, Milana Krush, Zhaohui Gu, Marsilius Mues, Philippe Depeille, Charles Mullighan, Jeroen P. Roose

## Abstract

Oncogenic mutations in *RAS* genes, like *KRAS^G12D^* or *NRAS^G12D^*, trap Ras in the active state and cause myeloproliferative disorder and T cell leukemia (T-ALL) when induced in the bone marrow via *Mx1CRE*. The RAS exchange factor RASGRP1 is frequently overexpressed in T-ALL patients. In T-ALL cell lines overexpression of RASGRP1 increases flux through the RASGTP/RasGDP cycle. Here we expanded *RASGRP1* expression surveys in pediatric T-ALL and generated a *RoLoRiG* mouse model crossed to *Mx1CRE* to determine the consequences of induced RASGRP1 overexpression in primary hematopoietic cells. RASGRP1-overexpressing, GFP-positive cells outcompeted wild type cells and dominated the peripheral blood compartment over time. RASGRP1 overexpression bestows gain-of-function colony formation properties to bone marrow progenitors in medium containing limited growth factors. RASGRP1 overexpression enhances baseline mTOR-S6 signaling in the bone marrow, but not *in vitro* cytokine-induced signals. In agreement with these mechanistic findings, hRASGRP1-ires-EGFP enhances fitness of stem- and progenitor- cells, but only in the context of native hematopoiesis. RASGRP1 overexpression is distinct from *KRAS^G12D^* or *NRAS^G12D^*, does not cause acute leukemia on its own, and leukemia virus insertion frequencies predict that RASGRP1 overexpression can effectively cooperate with lesions in many other genes to cause acute T cell leukemia.

## INTRODUCTION

Acute lymphoblastic leukemia (ALL) is an aggressive bone marrow (BM) malignancy. Approximately 15% of pediatric and 25% of adult cases of ALL are diagnosed as T-cell ALL (T-ALL) cog. In the clinic, T-ALL patients receive combination chemotherapy, sometimes combined with BM transplantation^2^. Understanding the molecularly altered biochemical pathways implicated in T-ALL is critical to developing effective molecular therapy (precision medicine). One commonly affected pathway is the RAS pathway with roughly 50% of all patient T-ALL cases demonstrating hyperactive RAS signals in the BM^3^. There are two major mechanisms of leukemogenic RAS signals in T-ALL patients; the first consists of somatic activating mutations of small RAS GTPases themselves and a second mechanism is through deregulated overexpression of the RAS activator *RASGRP1* (RAS guanine nucleotide releasing protein 1)^4,5^.

Oncogenic mutations in *RAS* are among the most common somatic mutations in cancer. Mutations in glycine at codon 12 in *KRAS* (*KRAS^G12^*) are prevalent in cancer and result in severely impaired GTPase activity and elevated levels of constitutively active, GTP-bound KRAS^6^. In hematopoietic malignancies, mutations in *NRAS* are 2 to 3 times more frequent than those in *KRAS*^7,8^, including pediatric T-ALL^9^. Generation of *Kras* mice aided the field of cancer research and oncogenic RAS signaling; these mice express a mutation of glycine to aspartic acid at codon 12 from the endogenous *Kras* locus in a controlled and inducible manner via a LoxP-STOP-LoxP cassette^6^. In BM cells, oncogenic KRAS can be inducibly expressed using *Mx1CRE* transgenic mice; in this model CRE is expressed from the IFN-α/β-inducible *Mx1* promoter by administration of polyinosinic-polycytidylic acid (pIpC)^10^. Such KRAS^G12D^ mice develop a lethal myeloproliferative disease (MPD) resulting in death around 35 days^11,l2^. In the background a T-ALL exists, which is suppressed by the MPD, but can be revealed via transplantation of KRAS^G12D^ hematopoietic stem cells into irradiated recipient mice^12–14^. *Mx1CRE-driven* NRAS^G12D^ mutation does not lead to acute MPD^15^. Instead mice develop a chronic myeloproliferative disorder and succumb to hematological disease by 15 months on a C57Bl/6 x 129/Sv.jae background but displaying a median survival of 588 days on a C57Bl/6 background^16^.

We reported deregulated overexpression of the RAS guanine nucleotide exchange factor *RASGRP1* in T-ALL patients^4,5^. RASGRP1 has a growth promoting role in T-cell leukemia^4^ and skin cancer^17^. RASGRP1 overexpression through retroviral transduction or via transgenic expression in thymocytes can trigger a leukemic phenotype^18–20^, but to date no genetic animal model exists to overexpress RASGRP1 in the BM in a controlled and inducible manner. As a consequence, mechanistic insights into overexpression of this RASGEF in an *in vivo* mouse model are lacking. Here we characterized a new *RoLoRiG* mouse model that allows for pIpC-induced overexpression of RASGRP1 and tracing of these BM cells with an ires-EGFP cassette. We report that overexpression of RASGRP1 results in increased baseline signals, increased spontaneous colony formation *in vitro*, and increased stem- and progenitor-cell fitness *in vivo* in the native bone hematopoiesis setting without acute leukemia development.

## RESULTS

### Inducible overexpression of hRASGRP1: RoLoRiG mouse generation

The small GTPase RAS is activated through RAS guanine nucleotide exchange factors (RASGEFs) and deactivated by RASGAPs (RAS GTPase Activating Proteins)^21,22^. We previously analyzed Affymetric gene array data on 107 pediatric T-ALL patients treated on COG (Children Oncology Group) studies 9404 and AALL0434^23^. We reported a unique range from low to high expression of *RASGRP1* that was not seen for *RASGRP2, RASGRP3*, or *RASGRP4*, nor for the other RASGEFs *RASGRF1, RASGRF2, SOS1*, and *SOS2*^4^. Likewise, expression levels of ten RASGAP family members where fairly even over these 107 pediatric T-ALL patients^24^. Capitalizing on HTSeq^25^ and DESeq packages^26^ for gene expression normalization, we next analyzed *RASGRP1* mRNA expression levels in 265 pediatric (n=250, age < 18yrs) and young adult (n=15, age ≥18) T-ALL from the COG AALL0434 cohort^23^ and observed a 100-fold range in expression levels (**Figure 1A**). Using integrated genomic analysis, Liu et al.^9^ identified six subsets of T-ALL that are characterized by six distinct genomic nodes and also represent different T cell development stages (**Figure 1B**)^9^. Examination of these *LMO2/LYL1* -, *HOXA-, TLX3-, TLX1*-, *NKX2-1*-, and *TAL1-* subsets did not reveal a clear pattern for *RasGRP1* expression based on these nodes (**Figure 1C**). During normal T cell development, *RasGRP1* levels are low in early thymocyte progenitors (DN, double negative), increase significantly in DP cells (CD4^+^CD8^+^; double positive) and peak in SP (single positive) thymocytes to drop again in peripheral T cells^27,28^. Thus, the *RasGRP1* expression we observe in the six genomic nodes does not follow the physiological pattern seen for normal mouse T cell development.

**Figure 1:**
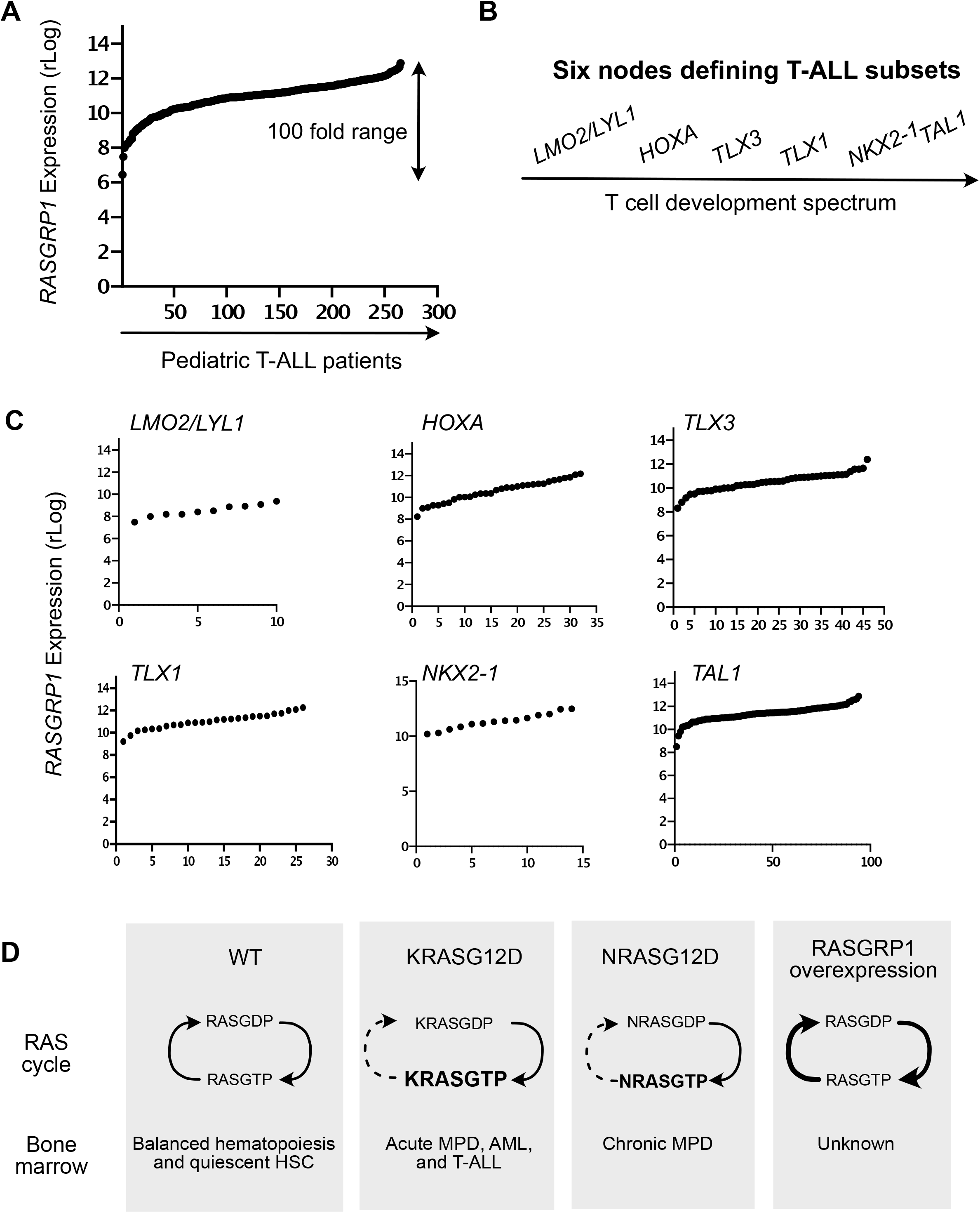
T-ALL patient analysis and mouse models. (A) Regularized log (rLog, normalized by DESeq2 R package) gene expression values of *RASGRP1* expression in 264 pediatric T-ALL patients. (B) Schematic of six subtypes in T-ALL. (C) rLog values of *RASGRP1* expression, plotted in the distinct *LMO2/LYL1* -, *HOXA-, TLX3-, TLX1*-, *NKX2-1*-, and *TAL1-* nodes. (D) Schematic representation of possible signaling networks in the BM of normal, oncogenic *Kras^G12D^* and *hRASGRP1* overexpressing mice.

Hematopoietic stem- and progenitor-cell homeostasis is regulated by the BM niche^29^ and cytokines released by stromal cells in this niche^30^. Cytokines can trigger RAS activation and RASGTP transmits signals to downstream effector kinase pathways, such as the RAF-MEK-ERK, Phosphatidylinositol 3-kinase (PI3K)-AKT and mTORC1-S6 and mTORC2-AKT pathways^5,21,31,32^. In T-ALL cell lines, KRAS^G12D^ causes high baseline RASGTP levels^5,33^, whereas overexpressed RASGRP1 constitutively loads RAS with GTP and RASGTP is constantly hydrolyzed back to inactive RASGDP^5^ (**Figure 1D**). *Kras^G12D^* in hematopoietic cells leads to elevated RASGTP levels at baseline^12,16^, whereas increases in RASGTP are more modest for *Nras^G12D^*^16^ Examination of RASGTP-effector kinase pathways in *Kras^G12D^* and *Nras^G12D^* BM subsets reveals differential perturbations depending on the cytokine stimulus used and the BM subset analyzed^15,16,34^. The biochemical and cellular consequences of RASGRP1 overexpression in BM are unknown (**Figure 1D**).

### A new RoLoRiG mouse model: Blood analysis reveals that hRASGRP1 overexpression bestows increased fitness to T-, B-, and myeloid-lineages

To create a model for overexpression of RASGRP1 in the BM, a cassette containing LoxP-STOP-LoxP, followed by hRASGRP1_-_ires_-_EGFP was knocked into the *Rosa26* locus^35^, in the reverse orientation and driven by the CAG promoter^36^. We termed this strain *RoLoRiG* (for **Ro**sa26-**Lo**xSTOPLox-**R**ASsGRP1-**i**res-**G**FP) (**Figure 2A**) and maintained it on a C57Bl/6 background. *RoLoRiG* mice were crossed to *Mx1CRE* mice and genotyped to verify *RoLoRiG* and *Mx1CRE* alleles (**Supplemental Figure S1A)**. A single dose of pIpC given to *RoLoRiG*^+/+^*Mx1CRE*^+^ mice resulted in expression of human RASGRP1 at levels in the range of what is observed for the human T-ALL cell line Jurkat^5,22^. In contrast, no hRASGRP1 expression was detected for *RoLoRiG^+/+^Mx1CRE* mice (**Figure 2B**).

**Figure 2:**
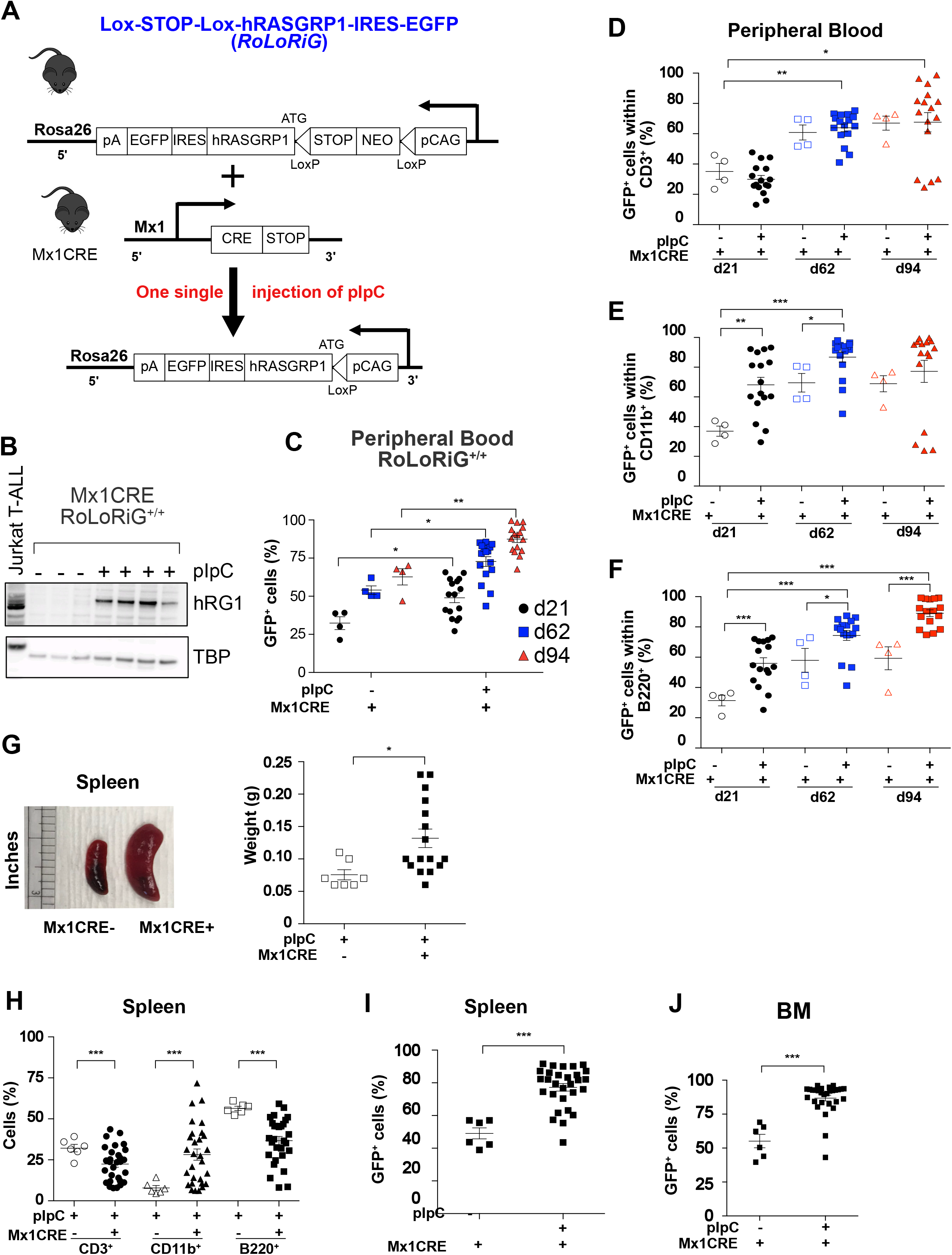
*In vivo* accumulation of *hRASGRP1*-overexpressing cells in *RoLoRiG* mouse model. (A) Schematic representation of the *RoLoRiG* model with overexpression of *hRASGRP1-*ires-EGFP. hRASGRP1 stands for human RASGRP1. *hRASGRP1-*ires-EGFP construct was knocked into the *Rosa26* locus. A stop codon is flanked by LoxP sites, which is excised when CRE recombinase is induced by injection of polyinosinic-polycytidylic acid (pIpC). (B) Representative Western Blot showing expression of hRASGRP1 detected with a human-specific monoclonal antibody in lymph node protein lysates of *RoLoRiG^+/+^ Mx1CRE^+^* mice, three months following pIpC injection. The Jurkat T-cell line was used as a positive control for hRASGRP1, TBP as loading control. (C) Percentages of GFP^+^ cells in peripheral blood of *RoLoRiG^+/+^ Mx1CRE^+^* mice on days 21, 62, and 94 after pIpC injection. Data are presented as mean ± SEM. Each symbol represents a single mouse in all figures (n>4 in Figure 2C). (D, E, F) GFP^+^ cells within of T-, myeloid-, and B-cells in *RoLoRiG*^+/+^*Mx1CRE*^+^ mice with or without pIpC injection on days 21, 62 and 94. (G) Splenic weights from *RoLoRiG^+/+^Mx1CRE* and *RoLoRiG*^+/+^*Mx1CRE*^+^ mice (n>7). Figures 2G–2J are at 94 days post pIpC injection. (H) Percentage of T-, myeloid-, and B-cells in spleens of *RoLoRiG^+/+^Mx1CRE*^−^ and *RoLoRiG*^+/+^*Mx1CRE*^+^ mice (n=6 or more). Increased percentages of CD11b^+^ cells were accompanied with decreases in B220^+^ B cells and CD3^+^ T cells. (I) Percentages of GFP^+^ cells in spleens of *RoLoRiG*^+/+^*Mx1CRE*^+^ mice with or without pIpC injection (n=6 or more). (J) Percentages of GFP^+^ cells in BM of *RoLoRiG*^+/+^*Mx1CRE*^+^ mice with or without pIpC injection (n>6).

We induced a cohort of *RoLoRiG*^+l+^ *Mx1CRE*^+^ mice (homozygous for the *RoLoRiG* allele,) with one single dose of pIpC and monitored health and general wellbeing. Overall, induced overexpression of RASGRP1 did not lead to a shortened life span or signs of severe lymphopenia and acute leukemia in a period of 400 days (data not shown). This is in sharp contrast to *Kras^G12D^Mx1CRE* mice, which develop a profound MPD to which mice succumb roughly one month after pIpC^11–14^, but similar to *Nras^G12D^Mx1CRE* mice on a C57Bl/6 background^16^. It should be mentioned that in the *Kras^G12D^Mx1CRE* studies mice were on a mixed background and several injections/doses of pIpC were administered to mice as opposed to one dose in our *RoLoRiG* study. Some *RoLoRiG* mice developed skin papillomas on their ears and tails, irrespective of pIpC injections and presumably caused by increased local IFN-α/β levels in the skin (**Supplemental Figure S1B**). Such skin papillomas are also reported in the *Kras^G12D^Mx1CRE* model^11^.

Tracking the mononuclear blood cell compartment longitudinally, we observed that GFP^+^ cells made up increasingly larger proportions as days 21, 62, and 94 evolve in *RoLoRiG*^+/+^*Mx1CRE*^+^ mice (**Figure 2C**) as well as in *RoLoRiG^+/-^Mx1CRE^+^* mice with one *RoLoRiG* allele (**Supplemental Figure S1C**). We observed modestly spontaneous increases in GFP+ cells without pIpC injection (**Figure 2C**), as the Mx1CRE cassette is known to cause a certain level of spontaneous CRE activity in most mouse facilities. Induced *Kras^G12D^ Mx1CRE* mice display severe MPD and increases in total white blood cells Analysis of T-, B- and myeloid-cells in peripheral blood from *RoLoRiG*^+/+^*Mx1CRE* mice revealed that hRASGRP1 overexpression does not lead to overt changes (**Supplemental Figures S2A-D**). As before, the *RoLoRiG^+/-^Mx1CRE* mice with only one *RoLoRiG* allele had a comparable phenotype (**Supplemental Figure S2E**) and for the remainder of the study we therefore limited assays to *RoLoRiG^+/+^Mx1CRE*^+^ mice and *RoLoRiG^+/+^Mx1CRE*-^−^ mice (without a functional CRE).

Notably, measuring the exact *<u>fraction</u>* of GFP^+^ cells, we observed that T-, B- and myeloid-cells with hRASGRP1-ires-EGFP expression efficiently outcompeted GFP^−^ cells in a time- and pIpC-dependent manner (**Figures 2D-F and Supplemental Figure S2F**). When pIpC is not administered, GFP^+^ cells eventually also outgrow negative cells because of the leakiness of *Mx1CRE*. At day 94, we observed mild splenomegaly in *RoLoRiG*^+/+^*Mx1CRE*^+^ mice (**Figure 2G**) and increased percentages of splenic CD11b^+^ myeloid cells (**Figure 2H**). The single, pIpC-driven genetic recombination event in BM cells resulted in a large proportion of GFP^+^ splenocytes over time (**Figure 2I**). Collectively, these results demonstrate that expression of RASGRP1-ires-GFP bestows an advantage over GFP^−^ cells and that this effect originates in the BM. Indeed, analysis of the BM from *RoLoRiG^+/+^Mx1CRE^+^* mice, which received one injection of pIpC showed enrichment for GFP^+^ cells (**Figure 2J**).

### hRasGRP1 overexpression does not impact homeostatic expansion following transplantation

Hematopoiesis and lineage development in the BM have classically been studied through transplantation of stem cells (LSK cells - Lineage^−^Sca^+^c-Kit^+^ cells) and allow for direct comparisons when done competitively. It should be noted that recipient mice receive irradiation to create space in the BM, resulting in an empty niche with abundantly available cytokines and many consider this an “injury and homeostatic expansion” model^37^. To assess lineage development by hRASGRP1-overexpressing hematopoietic stem cells in the BM, we introduced *RoLoRiG*^+/+^*Mx1CRE*^+^ or *RoLoRiG^+/+^Mx1CRE*^−^ LSK cells in a competitive manner (**Figure 3A**) using congenic marking through *CD45* alleles (**Supplemental Figure S3A**). *RoLoRiG^+/+^Mx1CRE*^−^ LSK cells were used as control to avoid the leakiness issue of *Mx1CRE*. CD45.2 *RoLoRiG^+/+^Mx1CRE*^+^ or *RoLoRiG*^+/+^*Mx1CRE*^−^ LSK cells filled the peripheral blood compartment with unexpected, similar efficiency and similar contribution to T-, B- and myeloid-cell lineages (**Supplemental Figures S3B-S3D**). Reconstitution by LSK cells with pIpC-induced hRASGRP1 overexpression resulted in modest splenomegaly (**Figure 3B**) and mildly increased percentages of CD11b^+^ myeloid cells in the spleen (**Figure 3C**).

**Figure 3:**
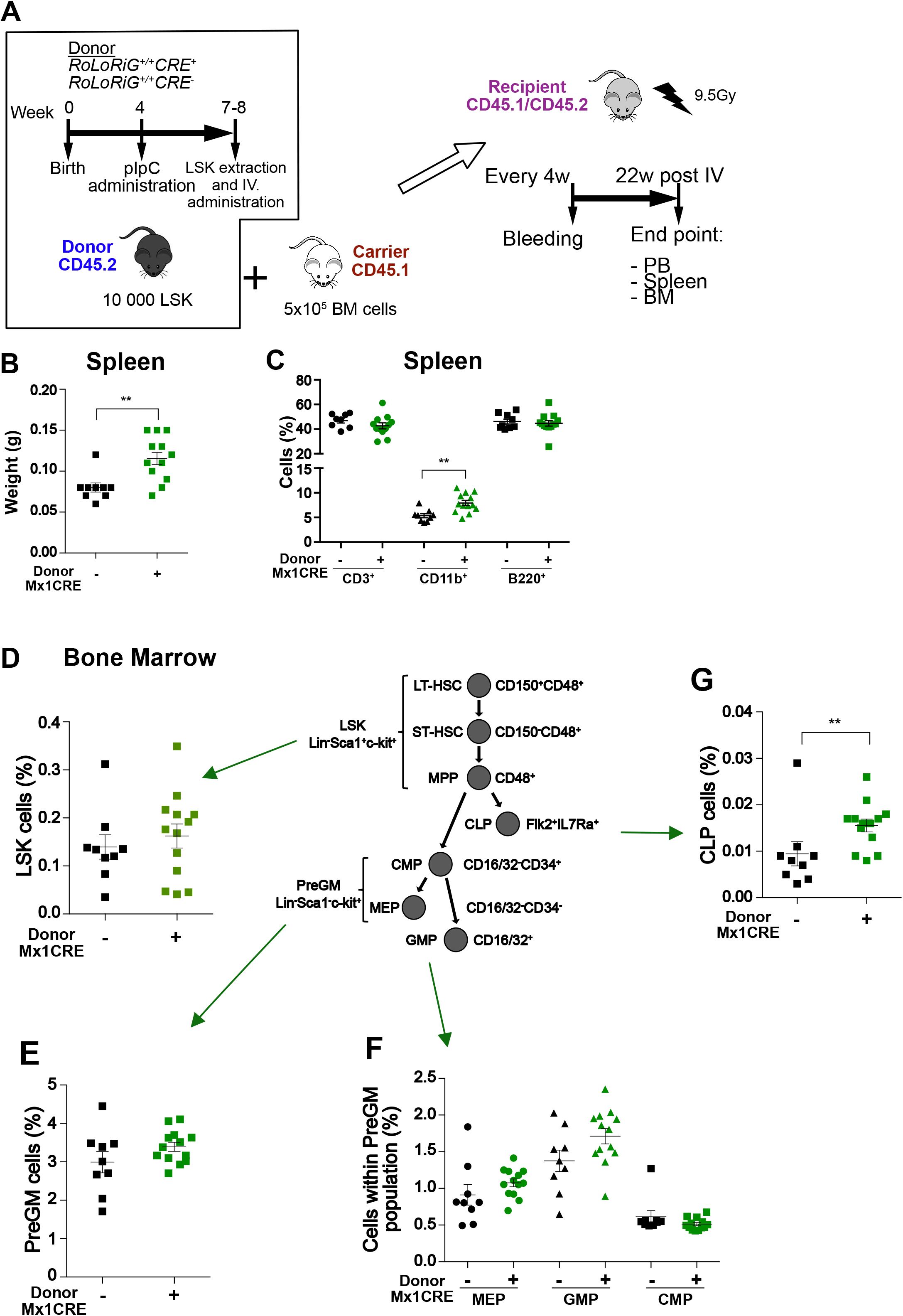
*hRASGRP1* overexpression in a bone marrow transfer model. (A) *RoLoRiG^+/+^Mx1CRE*^−^ or *RoLoRiG*^+/+^*Mx1CRE*^+^ mice were injected with pIpC at 4 weeks of age. BM cells were freshly isolated (7-8 weeks) from donor mice (CD45.2), which were either *RoLoRiG^+/+^Mx1CRE*^−^ or *RoLoRiG*^+/+^*Mx1CRE*^+^. Sorted 10 000 LSK cells were co-injected i.v. with 5×10^5^ carrier cells (CD45.1) into lethally irradiated recipients (CD45.1/CD45.2). Recipient mice were bled every 4 weeks until the final analysis at the endpoint (22 weeks). BM: bone marrow; LSK: Lin^−^Sca1^+^c-kit^+^; PB: Peripheral blood; pIpC: polyinosinic-polycytidylic acid. Data are presented as mean ± SEM of n=2 independent experiments. Congenic alleles allow for accurate enumeration of cell populations that are donor-derived (CD45.2) or are radiation-resistant host cells (CD45.1/CD45.2) (**Supplemental Figure S3A**). (B) Spleen weight from *RoLoRiG^+/+^Mx1CRE*^−^ and *RoLoRiG^+/+^Mx1* CRE^+^ mice at endpoint (n>9). Each dot represents a single mouse. **p<0.01. (C) Percentage of T-, myeloid and B-cells in spleen of *RoLoRiG^+/+^Mx1CRE*^−^ and *RoLoRiG*^+/+^*Mx1CRE*^+^ mice at endpoint (n>9). **p<0.01. (D-G) Analyses of percentages of LSK, PreGM, MEP, GMP and CMP within PreGM, and CLP in the BM. (n=9 or more). For additional subsets, see Supplemental Figure S4.

We next focused on the immature lineages in the BM (**Figures 3D-G** and **Supplemental Figures 4A and 4B**). In the context of this injury and homeostatic expansion model, *RoLoRiG^+l+^Mx1CRE^+^* transplanted LSK cells did not have an advantage compared to *RoLoRiG^+/+^Mx1CRE*^−^ LSK cells; we observed similar percentages of LSK, LT-HSC, ST-HSC, and MPP cell populations (**Figure 3E and Supplemental Figures S4A-S4E**). We also observed roughly equal numbers for PreGM, MEP, GMP, and CMP populations (**Figures 3FE and 3F**). CLP percentages were increased by two-fold in when *RoLoRiG^+/+^ Mx1CRE^+^* LSK cells are transplanted in comparison to *RoLoRiG^+l+^ Mx1CRE^−^* LSK **(Figure 3G)** and B220^+^ B-cells demonstrate a modest decrease (**Supplemental Figure S4F**). Thus, hRASGRP1-ires-GFP expressing LSK cells do not outperform wildtype LSK cells when investigated in a radiation/transplantation approach, which was somewhat unexpected given the fact that hRASGRP1-ires-GFP outgrow GFP-negative cell populations in a non-transplantation setting (**Figures 2 and 3A**). hRASGRP1-ires-GFP expressing LSK cells also contrast KRAS^G12D^ and NRAS^G12D^ expressing LSK cells, which are reported to display higher repopulating ability into the different cell lineages in the blood when transplanted into lethally irradiated mice^14,38^.

### Bone marrow characteristics under abundant cytokine stimulation

We previously uncovered that overexpression of RASGRP1 synergizes with cytokine receptor signals to aberrantly induce downstream RAS- and effector kinase-signals in T-ALL cell line models^4,5,22,39^, but these RASGRP1 signals have never been investigated in primary BM cells. Searching for mechanistic insights into the consequences of *RoLoRiG* in hematopoietic cells, we employed colony formation assays to determine the clonogenic capacity of hematopoietic progenitors as well as signaling assays (**Figure 4A**).

**Figure 4:**
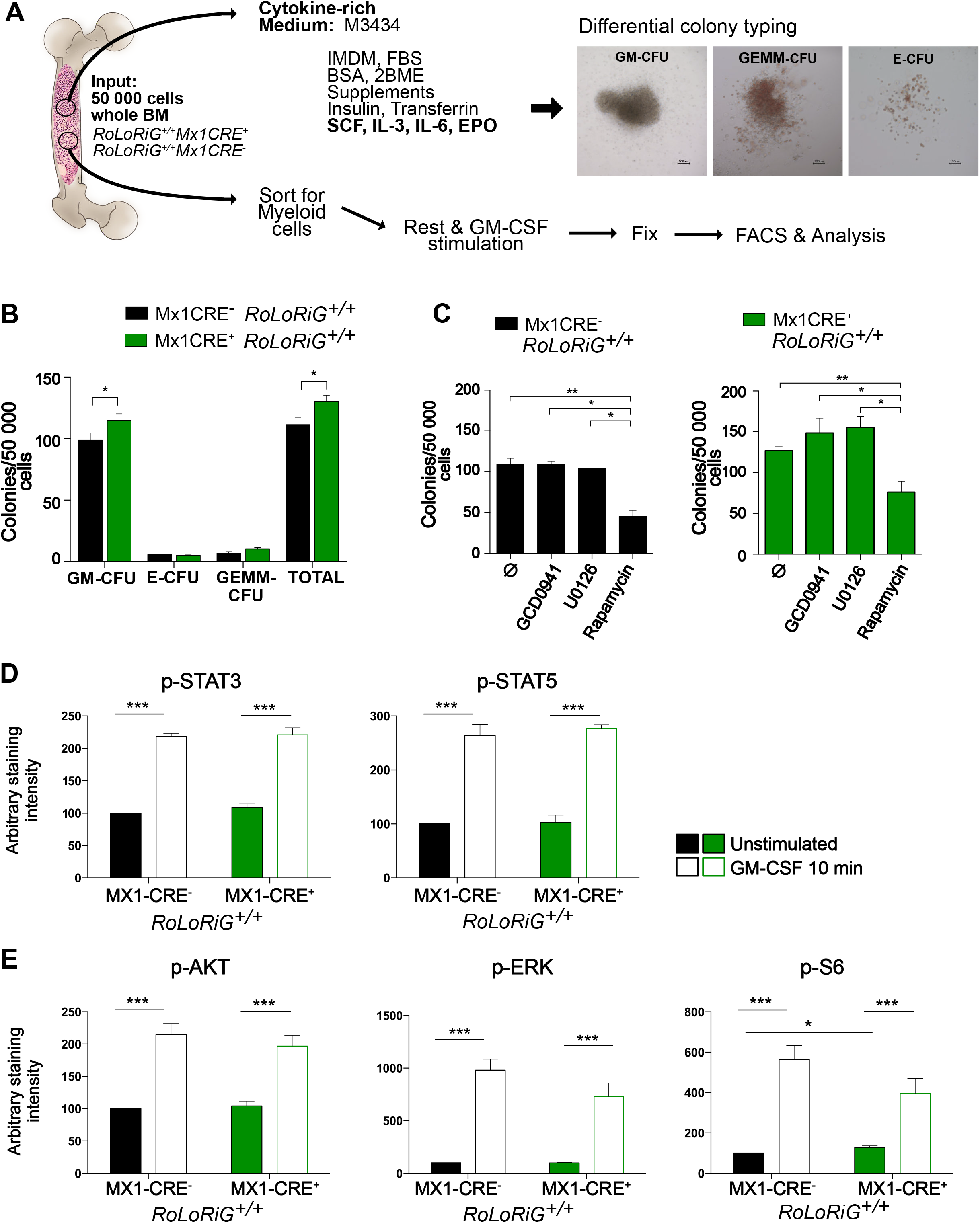
Cytokine-induced bone marrow colony formation and signaling. (A) 50 000 total BM cells from either *RoLoRiG^+/+^Mx1CRE*^−^ or *RoLoRiG*^+/+^*Mx1CRE*^+^ mice were seeded in cytokine-rich (M3434) medium. Representative images of the different colonies are shown after 7 days of incubation (100 μm Scale bar). Original magnification 10X. CFU: clonogenic forming units. For cytokine-induced signaling assays, cells were sorted, rested, stimulated, and processed for phosphor-antibody staining. (B-C) CFU potential of *RoLoRiG^+/+^Mx1CRE*^−^ and *RoLoRiG^+/+^Mx1CRE*^+^ cells grown in cytokine-rich medium (n>15). (B) Clonogenic (CFU) potential of *RoLoRiG^+/+^Mx1CRE*^−^ and *RoLoRiG*^+/+^*Mx1CRE*^+^ cells grown in cytokine-rich medium (n>15). (C) As in Figure 4B but in the presence of inhibitors of Ras kinase effector pathways: GDC0941 (0.1μM), U0126 (5μM) and Rapamycin (0.1μM) (n>3). (D) Analysis of p-STAT3 (Y705) and p-STAT5 (Y694) levels in rested and GM-CSF (40ng/ml) stimulated, sorted CD11B^+^ cells from *RoLoRiG^+/+^Mx1CRE*^−^ and *RoLoRiG*^+/+^*Mx1CRE*^+^ mice. N = 4 independent experiments. ***p<0.001. Unstimulated *RoLoRiG^+/+^Mx1CRE*^−^ samples were arbitrarily set at 100. For isotype control antibody staining see Supplemental Figure S5. (E) As in Figure 4D, assessment of p-AKT (S473), p-ERK (T202/Y204), and p-S6 (S235/S236) levels. N = 4 independent experiments. ***p<0.001, *p<0.05.

We explored clonogenic capacity of *RoLoRiG^+/+^ Mx1CRE^+^* or *RoLoRiG^+/+^Mx1CRE*^−^ BM cells with *in vitro* CFU (colony forming unit) assays in cytokine-supplemented M3434 CFU assay medium through differential quantification of the colonies based on morphological characteristics^40^. In cytokine-rich M3434 medium, overexpression of hRASGRP1-ires-EGFP resulted in modestly higher clonogenic capacity of total CFU and GM-CFU (**Figure 4B**). hRASGRP1-ires-EGFP did not impact the efficiency for E- and GEMM-CFU in M3434 medium (**Figure 4B**). RASGTP (active RAS) can signal to (PI3K)-AKT, AKT-mTOR (mammalian target of Rapamycin), and RAF-MEK-ERK resulting in stimulation of survival-, metabolic-, and cell cycle-changes^41,42^. Inclusion of pan-PI3K inhibitor GDC0941, MEK inhibitor U0126, and mTOR inhibitor Rapamycin revealed that the GM-CFU colony-forming capacity in cytokine-rich M3434 medium was only sensitive to addition of Rapamycin (**Figure 4C**). The effects of mTOR inhibition on colony formation in M3434 were both measurable in *RoLoRiG^+/+^Mx1CRE^+^* or *RoLoRiG^+/+^Mx1CRE^−^* BM cells (**Figure 4C**).

To investigate cytokine-mediated signaling in BM cells, we focused on myeloid cells given our GM-CFU results. Sorted CD11b^+^ *RoLoRiG^+/+^Mx1CRE* or *RoLoRiG^+/+^ Mx1CRE^+^* cells were rested and subsequently stimulated with GM-CSF (**Figure 4A**). GM-CSF stimulation resulted in the expected phosphorylation of STAT3 and STAT5 in either *RoLoRiG^+/+^Mx1CRE*^−^ or *RoLoRiG^+l+^ Mx1CRE*^+^ cells, indicative of efficient GM-CSF-JAK-STAT signaling (**Figure 4D and Supplemental Figure S5**). Downstream of RAS, GM-CSF-triggered modest induction of pAKT, but much more robustly induced pERK- and pS6-levels (**Figure 4E and Supplemental Figure S5**). In T-ALL cell lines, cytokines robustly trigger ERK and AKT pathways that are significantly reduced when RASGRP1-expression levels are decreased^4,39^. Surprisingly, hRASGRP1 expression did not impact the magnitude of the induction of these effector kinase pathways in sorted CD11b^+^ cells stimulated with GM-CSF. In fact, the only noticeable difference was the modestly increased p-S6 signal when comparing unstimulated *Mx1CRE*^−^ to *Mx1CRE^+^ RoLoRiG^+/+^* CD11b^+^ cells (**Figure 4E**).

### hRasGRP1 overexpression triggers spontaneous colony formation and baseline S6 signaling

Cytokines are produced and released by stromal cells residing in the BM niche and act on hematopoietic progenitors^30^, but stem cell niches are generally competitive environments^43^. To mimic a physiological *in vivo* situation of competition for limited amounts of available cytokines in the BM niche, we exploited cytokine-bare medium M3231^12^. M3231 was supplemented with low concentrations of either GM-CSF (0.01ng/ml) or of lymphoid-trophic IL-7 (0.1-10 ng/ml) (**Figure 5A**). As expected, control *RoLoRiG^+/+^ Mx1CRE*^−^ BM cells resulted in almost no colony formation under these conditions (**Figure 5B and 5C**). By contrast, *RoLoRiG*^+/+^*Mx1CRE*^+^ cells yielded substantial colony formation, both for GM-CSF and for IL-7 (**Figure 5B and 5C**). GDC0941, Rapamycin, and U0126 inhibitors all impacted the numbers of colonies by GM-CSF in M3231 **(Figure 5B**), implicating that PI3K-, mTOR-, and MEK-effector kinase signals all contribute to sustain colony formation and outgrowth in limiting growth factor conditions.

**Figure 5:**
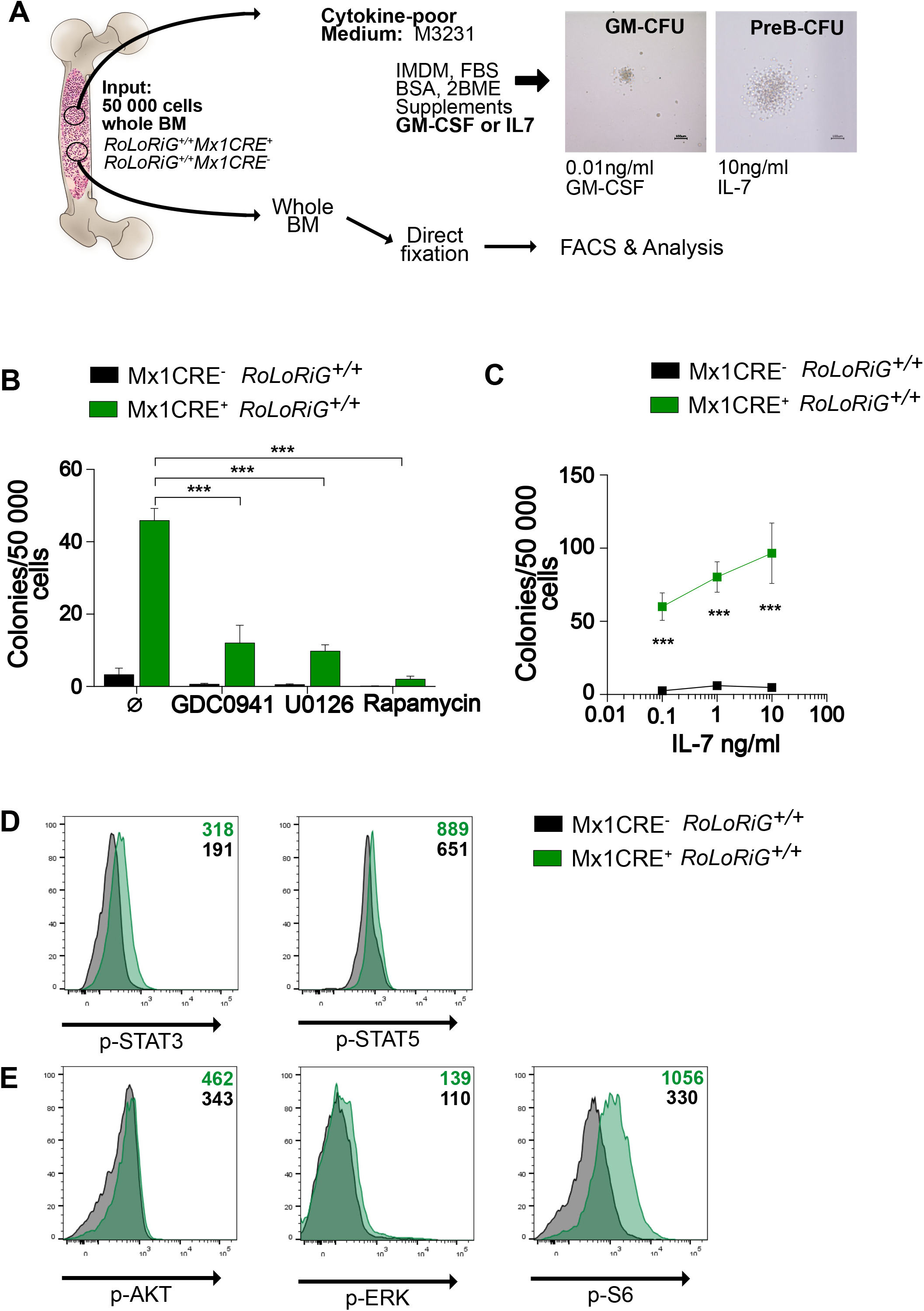
RasGRP1 overexpression drives spontaneous bone marrow colony formation and S6 signaling. (A) BM CFU analysis in cytokine-poor (M3231) medium. supplemented with either GM-CSF (GM-CFU) or IL-7 (PreB-CFU). Representative images following 7 days of incubation (100 μm Scale bar). Original magnification 10X. Schematic of baseline signaling assays in whole BM. (B) Clonogenic (CFU) potential of *RoLoRiG^+/+^Mx1CRE*^−^ and *RoLoRiG*^+/+^*Mx1CRE*^+^ cells grown in M3231 with 0.01 ng/ml of GM-CSF and inhibitors: GDC0941 (0.1μM), U0126 (5μM) and Rapamycin (0.1μM) (n>4). (C) Clonogenic (PreB-CFU) potential of *RoLoRiG^+/+^Mx1CRE*^−^ and *RoLoRiG*^+/+^*Mx1CRE*^+^ cells using 0.1, 1 and 10 ng/ml of IL-7 (n>3). (D) Analysis of baseline p-STAT3 (Y705), p-STAT5 (Y694), p-AKT (S473), p-ERK (T202/Y204), and p-S6 (S235/S236) signals in *RoLoRiG^+/+^Mx1CRE*^−^ and *RoLoRiG*^+/+^*Mx1CRE*^+^ total BM, immediately fixed after isolation. Mean fluorescence intensity is depicted by the numerical value. Data is representative of four independent experiments.

For the analysis of tonic signaling^44^ one isolates cells as quickly as possible followed by immediate fixation and analysis of cells *ex vivo* (**Figure 5A**). Analysis of tonic signals in freshly harvested total BM revealed that baseline pSTAT3- and pSTAT5-signals can be detected when the staining of cells with phospho-specific antibodies is compared to staining with isotype control antibodies (**Figure 5D**). These baseline pSTAT3- and pSTAT5-signals likely reflect the cytokine- and chemokine-signals received by BM cells immediately prior to harvesting from their niche and immediate fixation. Measurements of baseline pAKT-, pERK-, and pS6-levels that lie downstream of RAS revealed that activities through AKT- and S6-pathways in hematopoietic cells were more prevalent on a whole than the ones through the ERK pathway (**Figure 5E**). Moreover, when comparing *RoLoRiG^+/+^ Mx1CRE^+^* BM to *RoLoRiG^+/+^ Mx1CRE*^−^ BM, we observed increased baseline S6 signals when hRASGRP1 is overexpressed (**Figure 5E**).

### hRASGRP1 overexpression results in increased fitness of stem cells in the native BM

In general, a classic hematopoietic developmental scheme is believed to follow a hierarchical order, as depicted in **Figure 3**^45,46^. However, important nuances have been uncovered when native hematopoiesis is compared to the hematopoiesis that occurs following irradiation and transplantation^37,47^. Focusing on native hematopoiesis, *RoLoRiG*^+/+^*Mx1CRE*^+^ mice receiving one dose of pIpC revealed significantly increased percentages in LSK cells (**Figure 6A**) with increased LT-HSC but comparable ST-HSC (**Figure 6B**). In addition, higher percentages of MPP (**Figures 6C**) and CLP (**Figures 6D**) were detected in *RoLoRiG*^+/+^*Mx1CRE*^+^ mice. Increases in PreGM cells (**Figure 6E**) stemmed from higher percentages of MEP and GMP in *RoLoRiG^+/+^ Mx1CRE^+^* mice with normal CMP (**Figure 6F**).

**Figure 6:**
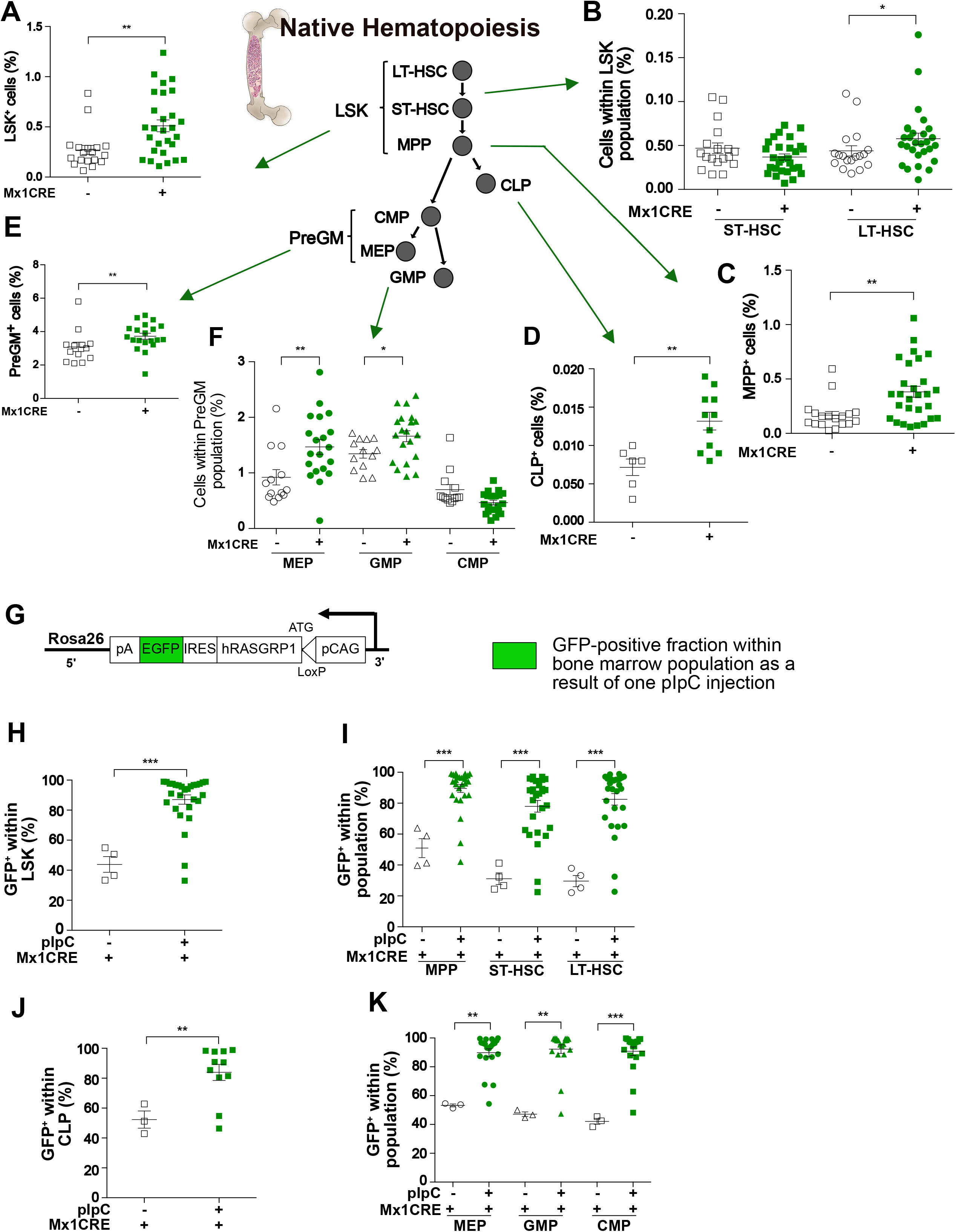
Increased fitness in Native hematopoiesis in RoLoRiG^+/+^ mice. (A) Percentage of LSK in the BM of *RoLoRiG^+/+^Mx1CRE*^−^ or *RoLoRiG^−/+^Mx1CRE^+^* mice (n>18). In panel 6A and all subsequent panels in figure 6, data are presented as mean ± SEM. Each symbol represents a single mouse. *p<0.05, **p<0.01, ***p<0.001. (B-F) Percentage of specific developmental stages in the BM *RoLoRiG*^+/+^*Mx1CRE*^−^ or *RoLoRiG*^+/+^*Mx1CRE*^+^ mice, analogous to the developmental gating scheme used in Figures 3. (G) Inclusion of the ires-EGFP cassette allows for specific assessment of the GFP^+^ fraction within each of the hematopoietic subsets within the BM. (H-K) Proportions of GFP^+^ cells within in *RoLoRiG*^+/+^*Mx1CRE*^+^ mice with or without pIpC injection. (H. LSK; I. MPP, ST-HSC and LT-HSC cells; J. CLP cells; K. MEP, GMP and CMP cells.

The inclusion of the ires-EGFP cassette allowed for quantitative assessment of the fraction of GFP^+^ cells in each of the BM stem- and progenitor-populations, examined 94 days after one single dose of pIpC (**Figure 6G**). These analyses revealed a striking pattern: in *RoLoRiG^+l+^Mx1CRE^+^* mice all populations revealed percentages of GFP^+^ cells that approached 100 percent (**Figures 6H–6K**). By contrast, in *Kras^G12D^Mx1CRE* mice, BM FLK2^−^LSK cell numbers are reduced by 2-fold following 2 weeks of pIpC injection, while numbers of these cells increase in spleen^14^. Thus, the one-time induced GFP^+^ fraction takes over the BM in a sustained manner, demonstrating that RASGRP1 overexpression provides increased fitness to the stem cell populations.

## DISCUSSION

High levels of *RASGRP1* expression are seen in many T-ALL patients and the *RoLoRiG* model allowed for the analysis of signaling- and functional-traits as a result of hRASGRP1 overexpression in primary hematopoietic cells for the first time. We established that hRASGRP1 overexpression bestows increased fitness onto stem- and progenitor-cells without causing acute leukemia, a phenotype that is very distinct from oncogenic KRAS^G12D^ expression driven by Mx1CRE and more similar to NRAS^G12D^ expression in the BM.

The fact that hRASGRP1-ires-EGFP or NRAS^G12D^ expression^15,16,34,38^ in the BM alone do not lead to acute leukemia, the notion that *NRAS* mutations are more frequent than *KRAS* mutations in leukemia^7–9^, and our reports that overexpression of RASGRP1 is often observed in T-ALL patients^4^, suggests that additional hits play a role. This “two-hit” theory was already described almost 50 years ago by Knudson *et al*.^48^, many research groups have based their work on this theory^49–52^, and Molony leukemia virus insertions in genes such as *Evi1* in the NRAS^G12D^ mouse result in acute leukemia^16^. To explore “two-hit” for RASGRP1 overexpression we plotted co-occurrence for the top 8 SL3-3 leukemia virus insertion sites in our previously reported mouse leukemia screen^4^. Such analyses can predict redundancy and cooperation in oncogenesis. For example, the *Pvt1* locus was demonstrated to encode microRNAs that control *MYC* expression and co-insertions for *Pvt1* and *MYC* are not found in these SL3-3 driven leukemias^53^ (**Figure 7**). SL3-3 insertions in *RasGRP1* frequently co-occur with insertions in *Evi5, Notch1, Ahi1, GFi1, Myc, RRas2*, and particularly *Pvt1* (**Figure 7**) and future work is required to understand the mechanisms here. For example, does the increased fitness from RASGRP1 expression allow for acquisition of additional mutations without decreasing viability of the (pre) leukemic cell or does RASGRP1 have bimodal functions such as the ones described for NRAS^G12D^ with stem cell renewing- and proliferation enhancing-properties in distinct cell subsets^38^?

**Figure 7:**
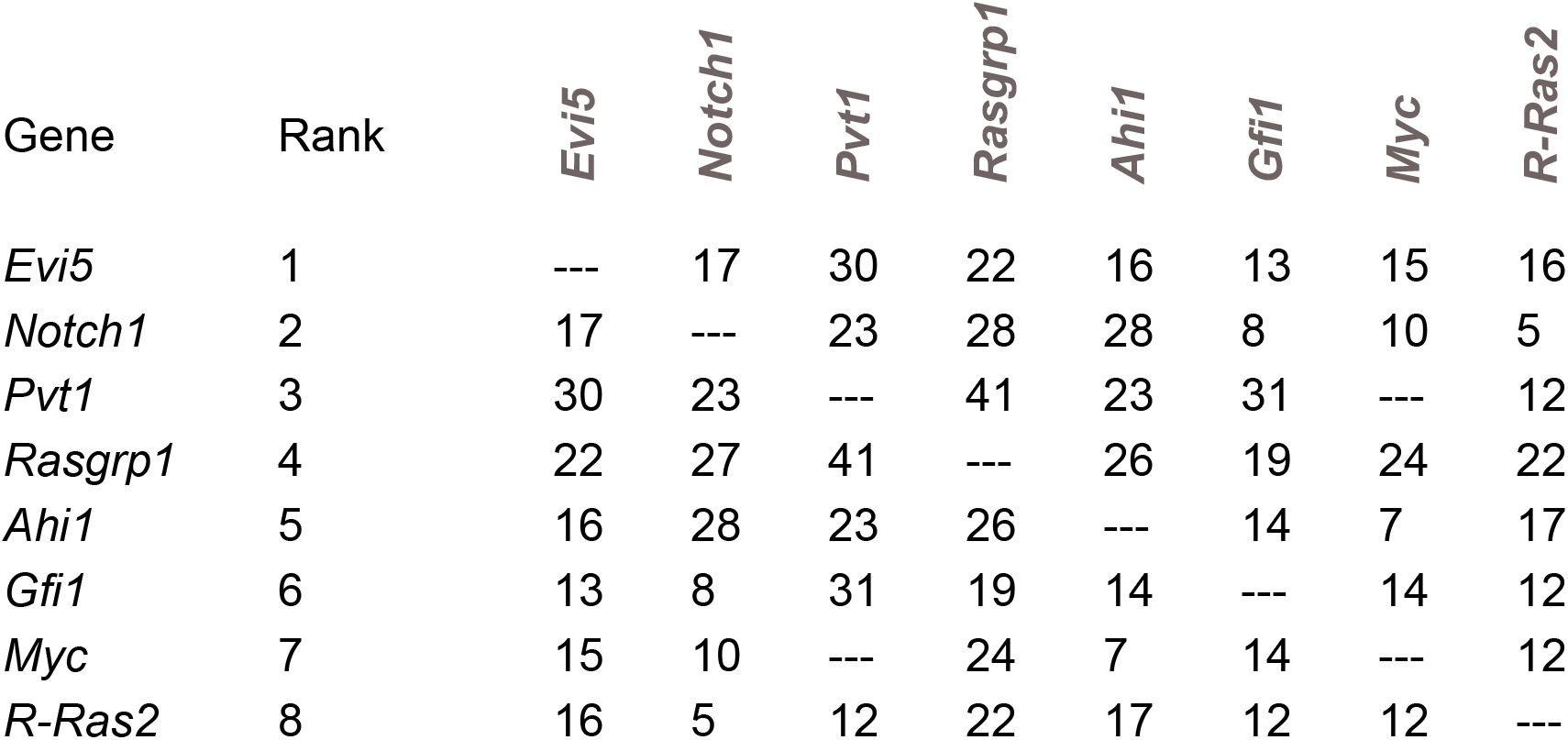
Co-insertion table of SL3-3 leukemia virus insertions. SL3-3 leukemia virus insertions that resulted in T-ALL in mice were mapped against each other to understand the relationship between the top 8 SL3-3 insertion sites.

Prominent features in our *RoLoRiG* model are the increased stem cell fitness in native hematopoiesis, the increased (mTOR) S6 signals at baseline, and the enhanced spontaneous colony formation. mTOR is an interesting node in hematopoiesis and leukemia that deserves more research. TSC (Tuberous sclerosis) genes are negative regulators of mTOR signaling and downregulation of TSC2 gene expression has been reported in BM samples of acute leukemia patients^54^. Raptor is a critical component of the mTORC1 complex^32^ and Raptor deletion results in compromised *ex vivo* colony formation with small irregular colonies^55^, corroborating our results with the mTOR inhibitor Rapamycin. Deletion of Raptor by a Mx1CRE driver does not lead to altered BM progenitor cell frequencies However, in the context of T-ALL with *PTEN* deletion, Raptor deletion reduces T-ALL, arguing that mTORC1 is a critical effector in leukemia with *PTEN* loss^55,56^. Loss of Rictor, component of mTORC2, has little effect on normal HSC function, but similarly to Raptor deletion, deletion of Rictor normalizes the HSC proliferation in PTEN null mice and also prevented the T-ALL^56,57^. Baseline signaling has remained a relatively understudied area of signal transduction and we believe it will be of interest to carefully characterize baseline signals in the context of native hematopoiesis^37,47^ and leukemia.

## MATERIALS AND METHODS

### Plasmid cloning and generation of mice with hRasGRP1 over-expression

The expression construct was generated by cloning the various components into targeting ROSA vector (ROSA-HR). Vector was obtained by introducing first a flanked Neo/STOP cassette and then the *hRasGRP1* cDNA cassette, _IRES_-EGFP, and human polA sequences into a Rosa26 targeting vector that already contain a ubiquitous CAG promoter cassette (**Figure. 1A**). This construct was transfected into ES cells by electroporation in optimized condition (5×10^6^ cells in presence of 40 μg linearized plasmid, 260V, 500 uF). Positive selection started 48 h after electroporation by adding 200 μg/ml of G418 in 96-well plates. Positive clones were further screened by southern blotting. Furthermore, the hybridization with internal probe did not indicate that any of the ES cells screened contained an additional randomly integrated copy of the targeting plasmid. Mice received a single intraperitoneal injection of polyinosinic-polycytidylic acid (pIpC) (1μg/250μl/mouse) to induce the expression of hRASGRP1 and GFP from the *Rosa26* locus.

### Genotyping

See Supplemental Figure S1

### Western blotting

To analyze hRASGRP1 overexpression under basal conditions by western blot, cells of interest were washed with phosphate-buffered saline (PBS) and lysates were prepared by adding NP40 lysis buffer supplemented with protease and phosphatase inhibitors [10 mM sodium fluoride, 2 mM sodium orthovanadate, 0.5 mM EDTA, 2 mM phenylmethylsulfonyl fluoride, 1 mM sodium molybdate, aprotonin (10 mg/ml), leupeptin (10 mg/ml), pepstatin (1 mg/ml)]. Lysates were incubated for 20 min at 4°C, centrifuged at 13 000rpm and debris-free lysates were transferred to new collection tubes. Western blotting analysis was performed with antibodies specific for hRASGRP1 (H120, Santa Cruz Biotechnology, Dallas, TX, US) and TBP (#8515, Cell Signaling, Danvers, MA, US). Western blots were visualized and quantified with enhanced chemiluminescence and imaging on a Fuji LAS 4 000 image station (GE Healthcare, Chicago, IL, US)^4^.

### Single cell suspension generation from tissues

Mice were euthanized 3 to 6 months after pIpC injection. Tibia and femurs of mice were collected and cleaned from flesh. Bones were then crushed using mortar and pestle in Hank’s Balanced Salt Solution (HBSS; UCSF cell culture facility) containing 2% of FBS. Cells were filtered through a 70 μm strainer. Spleen and thymus were harvested from the mice and mashed through a 70 μm cell strainer. Red blood cell lysis was performed for all tissues with ACK buffer (150 mM NH_4_Cl, 10 mM KHCO_3_, 0.1 mM Na_2_EDTA at pH 7.2-7.4). Bone marrow (BM) cells were washed once with HBSS, then loaded onto a Histopaque-1119 (Sigma, San Luis, MO, US) density gradient and centrifuged at 1500 rpm with no break for 5 min to separate bone debris from cells. Splenocytes and thymocytes were washed once with HBSS/2% FBS. All cells were then counted and prepared for experimental setup.

### LSK cell transplantation

BM cells from *RoLoRiG^+/+^Mx1CRE*^−^ or *RoLoRiG*^+/+^*Mx1CRE*^+^ mice were isolated as described earlier. Cells were blocked with Rat IgG and stained for lineage markers (CD3, CD4, CD5, CD8, CD11b, B220, Ter119), c-Kit and Sca-1 under sterile conditions. LSK cells were then sorted using FACS Aria by gating on GFP^+^Lin^−^cKit^+^Sca1 ^+^ population in *RoLoRiG*^+/+^*Mx1CRE*^+^ samples or on all cells in *RoLoRiG^+/+^Mx1CRE*^−^ samples. CD45.1^+^/CD45.2^+^ mice were lethally irradiated (9.5Gy) and injected in the lateral tail vein^58^ with 10×10^3^ LSK from either *RoLoRiG^+/+^Mx1CRE*^−^ or *RoLoRiG*^+/+^*Mx1CRE*^+^ (CD45.2^+^) mice. Each LSK sample was mixed together with 5×10^5^ unfractionated adult BM cells (CD45.1^+^). Donor chimerism in peripheral blood (PB) was assessed by flow cytometry at 4, 6, 10, 14 and 18 weeks post transplantation. For hematopoietic tissue analysis, mice were euthanized at 18 and 22 weeks after cell transplantation.

### Hematopoietic and donor chimerism analysis

PB was collected from mice tail vain into a tube containing PBS with FACS buffer (2mM EDTA, 2% FBS and 0.09% NaN_3_). Blood cells were lysed using ACK buffer. Cells were centrifuged for 5 minutes at 500 g, supernatant was discarded, and cells were suspended in FACS buffer for flow cytometry staining. The following antibodies were used: α-CD3-APC, α-CD11b-VioletFluor450, α-B220-PerCP-Cy5.5, α-CD45.1-PE-Cy7, α-CD45.2-PE to analyze T-, B- and myeloid cells and the chimerism after LSK transplantation.

BM cells were blocked with Rat IgG for 30 minutes at 4°C. Cells were stained with lineage markers (CD3, CD4, CD5, CD8, CD11b, B220, Ter119); c-kit and Sca-1 for LSK and PreGM populations; CD150 and CD48 for MPP, ST-HSC and LT-HSC; CD16/CD32 and CD34 for CMP, GMP, MEP populations; Flk2 and IL7Rα for CLP population as previously described **(Fig 3A)**^59,60^. Samples were acquired on a BD LSRII Fortessa (BD, Franklin Lakes, NJ, US). FlowJo software was used to analyze the data. Antibodies information is on Table 1.

### Clonogenic forming unit assay

Primary mouse clonogenic progenitor assays or colony forming units (CFU) assays were performed by plating 5×10^5^ unfractionated nucleated BM cells in methylcellulose M3434 (StemCell Technologies, Vancouver, Canada) in 1 ml duplicates. Colonies were counted and scored after 7 days of incubation using standard morphological criteria^40^. To assay granulocyte/macrophage (GM-CFU), unfractionated nucleated BM cells (5×10^5^) were suspended in methylcellulose medium (M3231, StemCell Technologies) with 0.01 ng/ml murine GM-CSF (Peprotech, Rocky Hill, NJ, US) and plated in 1 ml duplicates. Colonies were counted after 7 days. To assay PreB-cell colonies (PreB-CFU), unfractionated nucleated BM cells (5×10^5^) were suspended in methylcellulose medium (M3231, StemCell Technologies) with 0.1, 1 and 10ng/ml murine IL-7 (Peprotech) and plated in 1-ml duplicates. Colonies were counted after 7 days. Representative images were taken with a Keyence BZ-X710 microscope (Keyence, Itasca, IL, US). Images were viewed with the Keyence BZ-X viewer and BZ-X analyzer. Image brightness and contrast were edited using Affinity Designer. To assess the effect of known chemical inhibitors on CFUs from either *RoLoRiG^+/+^Mx1CRE*^−^ or *RoLoRiG*^+/+^*Mx1CRE*^+^ mice, cells were plated in the absence or presence of U0126 MEK inhibitor (5μM), PI3K inhibitor GDC-0941 (0.1 μM) or mTOR inhibitor rapamycin (0.1 μM).

### Statistics

Results are shown as mean□±□SEM and the normality of data was checked using the D’Agostino-Pearson test. For variables with a normal distribution, Student’s t test and one-way ANOVA were used to compare the significance of differences between experimental groups. For variables with no normal distribution, Mann–Whitney U test or Kruskal–Wallis test was applied. Statistical analysis was performed using GraphPad Prism version 6.00. A p-value<0.05 was considered significant and indicated with an asterisk.

## ACKNOWLEDGEMENTS

This work was supported by an Alex’ Lemonade Stand Foundation Innovator Award, the NIH/NCI (R01 - CA187318), and the NIH/NHLBI (R01 - HL120724) (all to JPR). Further support came from a Leukemia & Lymphoma Society grant (to MM) and the Rothschild Fellowship for postdoctoral fellows in the Natural, Exact or Life Sciences and Engineering (to LK), and PD is a Mark Foundation Momentum Fellow supported by a fellowship from the Mark Foundation for Cancer Research; and by NCI grants CA021765 (St Jude Comprehensive Cancer Center Support Grant), an NCI R35 Outstanding Investigator Award (R35 CA197695) and a St. Baldrick’s Foundation Robert J. Arceci Innovation award. We thank the members of the Roose lab and the Heme-Onc community at UCSF for useful suggestions and comments. We thank UCSF flow cytometry facility and DRC Center Grant NIH P30 DK063720. We thank Emmanuelle Passague and her lab for kindly providing us *Mx1-CRE* mice.

## AUTHORSHIP

Contributions: L.K., D.R-M., O.K. and M.M. performed experiments and analyzed results. P.D. made the mouse construct. Z.G. and C.G.M. generated and analyzed human T-ALL genomic data. L.K and D.R-M. made the figures; J.R. designed the research and secured the majority of the funding. L.K., D.R-M. and J. R. wrote the paper.

## Current affiliations

The Current affiliation of O.K. is Liggins Institute, The University of Auckland, Auckland, New Zealand. The current affiliation of M.M. is Miltenyi Biotec GmbH, Friedrich-Ebert-Str. 68, 51429 Bergisch Gladbach, Germany.

