## Supplementary figures and images for "Increased baseline RASGRP1 signals Enhance Stem Cell Fitness during Native Hematopoiesis"

### Supplemental Figure 1

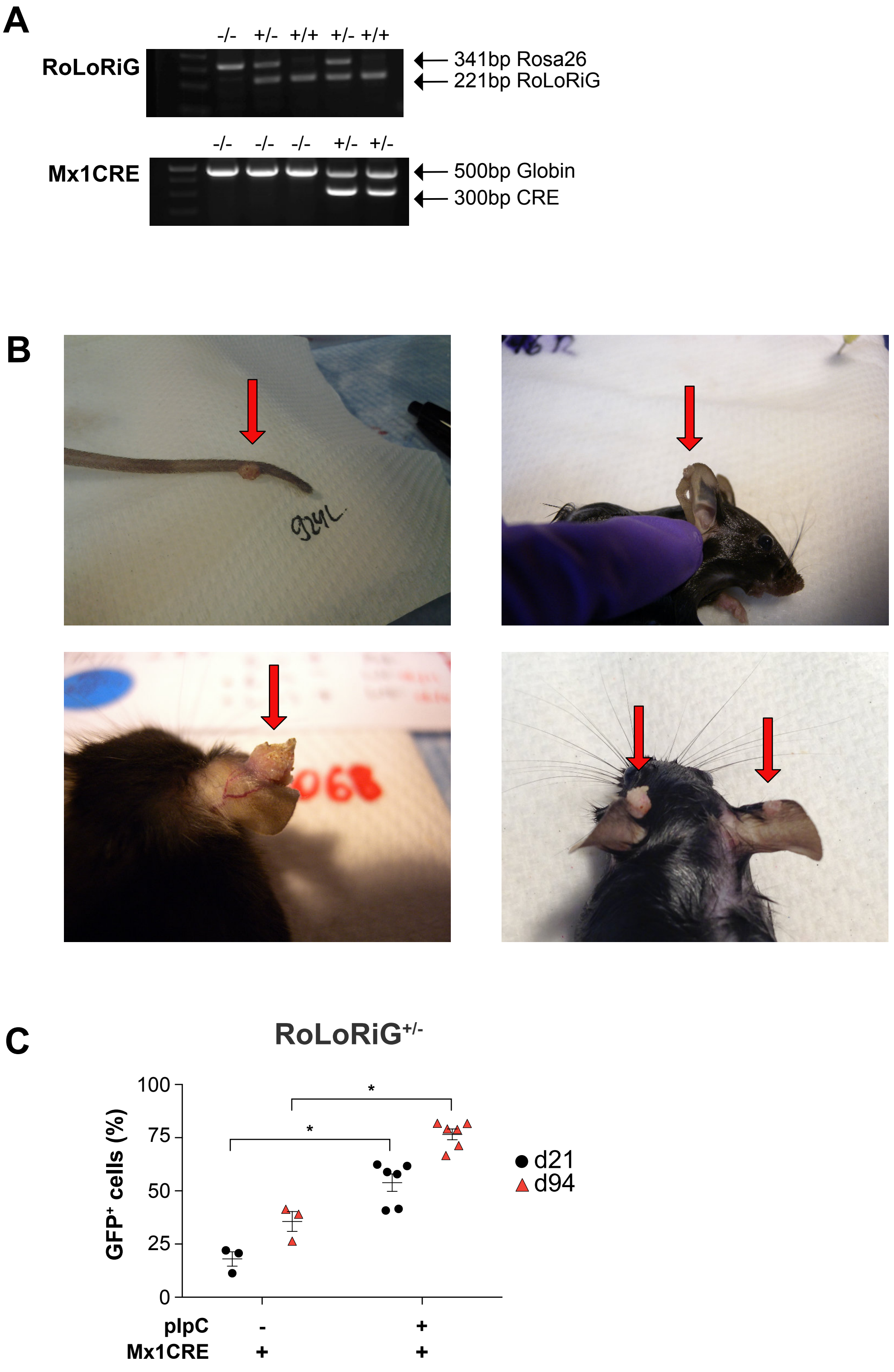


Karra, Romero-Moya, Ksionda et al.
Supplemental Figure S1

### Supplemental Figure 2

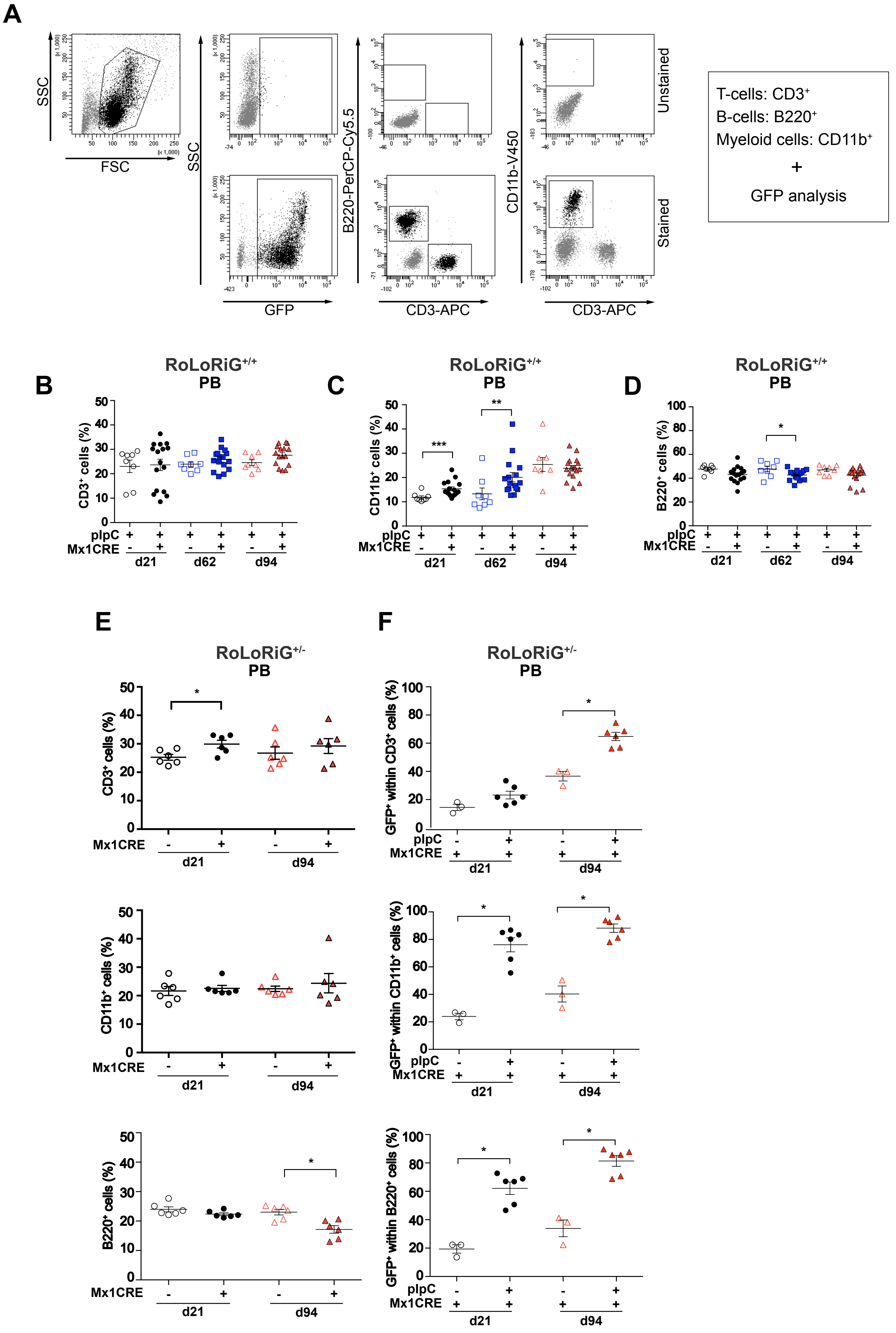


Karra, Romero-Moya, Ksionda et al.
Supplemental Figure S2

### Supplemental Figure 3

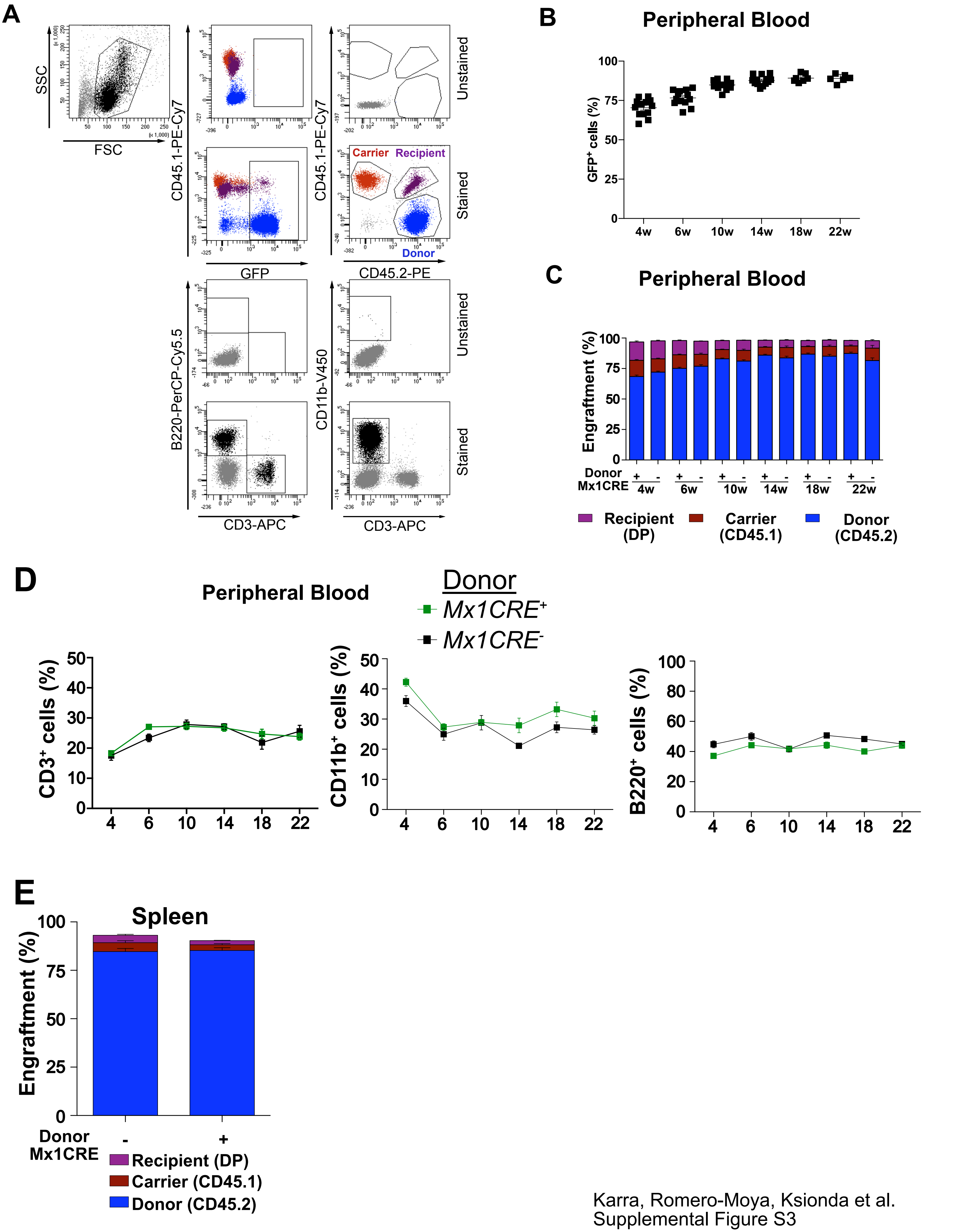


Karra, Romero-Moya, Ksionda et al.
Supplemental Figure S3

### Supplemental Figure 4

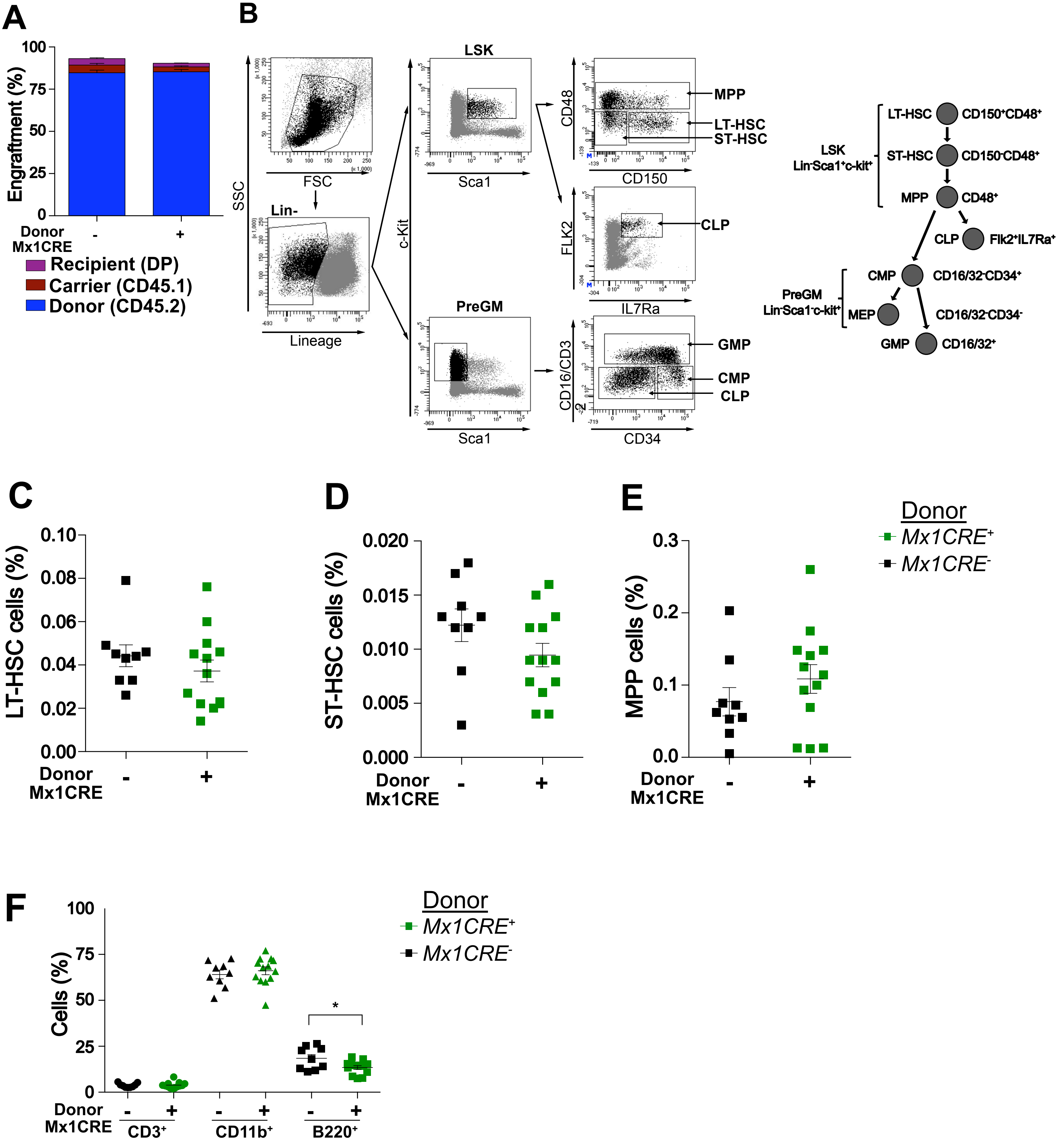


Karra, Romero-Moya, Ksionda et al.
Supplemental Figure S4
