## Supplemental Figure 5 for "Increased baseline RASGRP1 signals Enhance Stem Cell Fitness during Native Hematopoiesis"

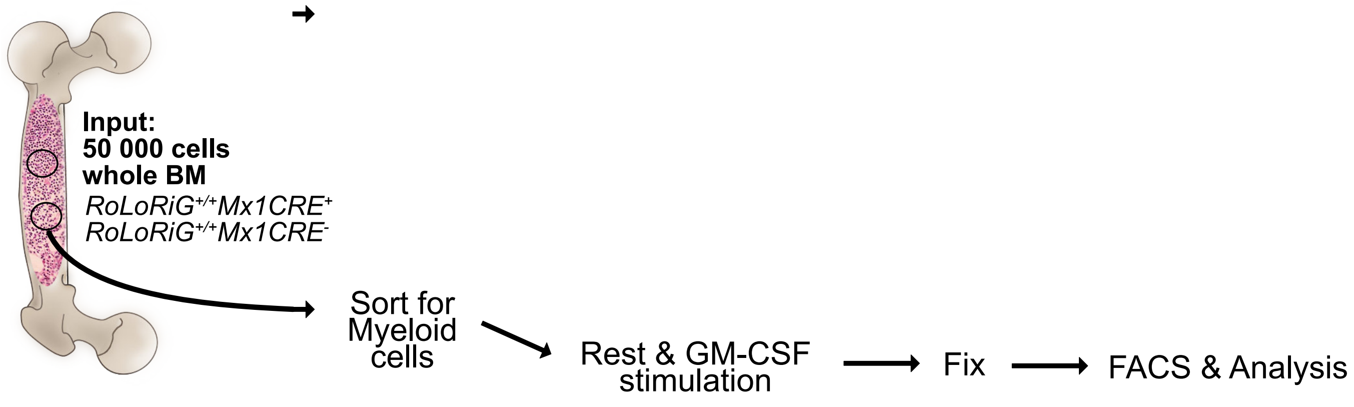


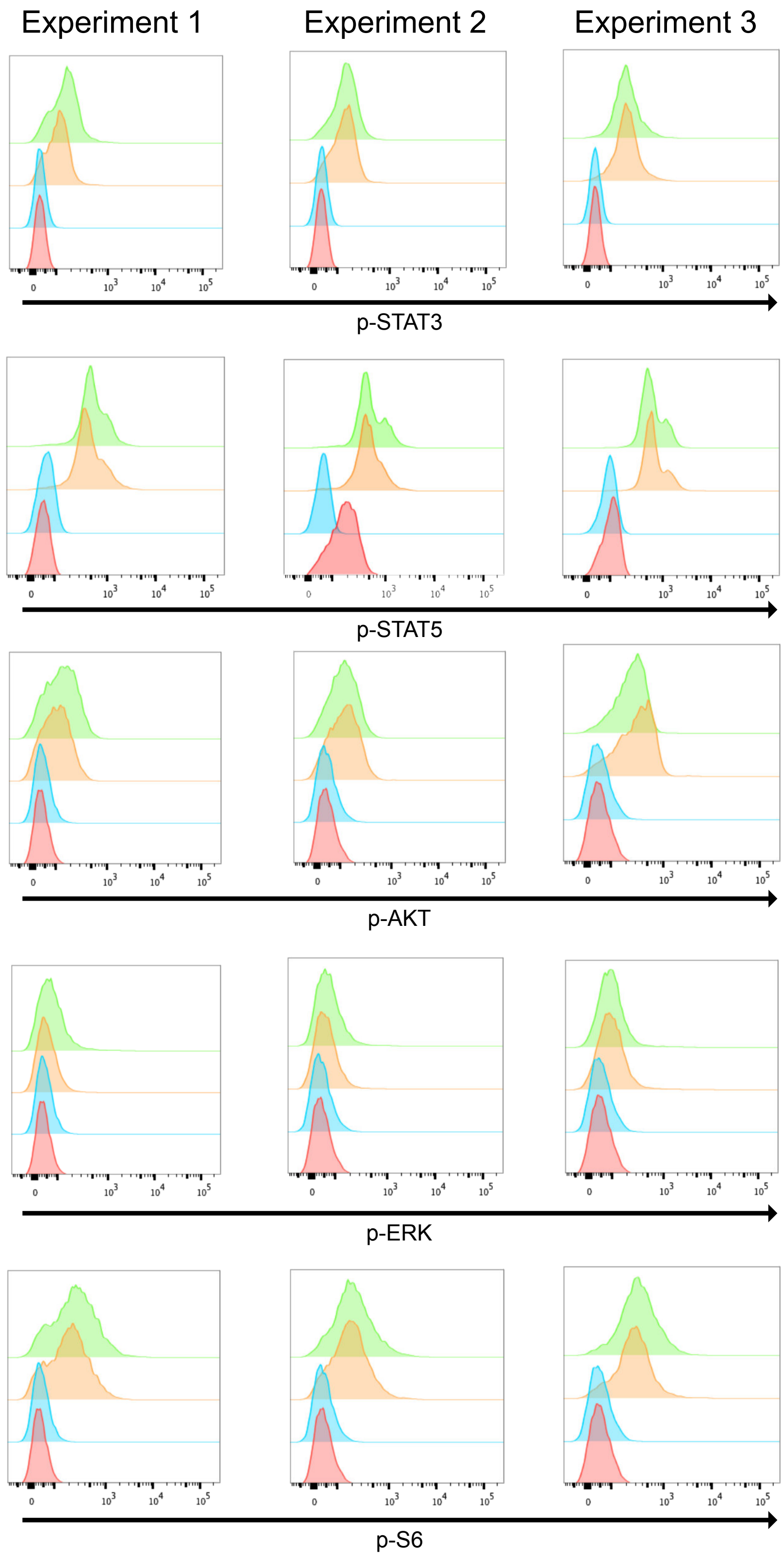


Histograms of phospho-stainings on **rested, unstimulated samples** (Figure 4D and 4E) to illustrate the differences between isotype control antibody staining and specific, phospho-antibody staining

- RoLoRiG$^{+/+}$MX1-CRE$^{+}$ Phospho-Ab stain
- RoLoRiG$^{+/+}$MX1-CRE$^{-}$ Phospho-Ab stain
- RoLoRiG$^{+/+}$MX1-CRE$^{+}$ Isotype control stain
- RoLoRiG$^{+/+}$MX1-CRE$^{-}$ Isotype control stain

Karra, Romero-Moya, Ksionda et al.
Supplemental Figure S5
