## Supplemental Figure legends for "Increased baseline RASGRP1 signals Enhance Stem Cell Fitness during Native Hematopoiesis"

**Karra et al., Legends Supplemental Figures S1-S5**

**Supplemental Figure S1**

(S1A) The unarranged Rosa26 is identified by a 341 bp product. Recombined *RoLoRiG* is identified by a 221bp product. The Mx1CRE allele is identified by a 300bp product. Globin was used as internal control (Ctrl). Genotyping was carried out on tails digested at 55 °C for 4-6 hours in a buffer containing 100mM Tris HCl (pH 8.5), 5 mM EDTA (pH 8), 0.2% SDS, 200 mM NaCl and 200 𝛍g/ml Proteinase K (Roche, #3115801). Samples were centrifuged at 13000 rpm, 5 min, supernatant was collected into clean tubes and DNA was precipitated with methanol at a 1:1 ratio. Genomic DNA was washed with 70% EtOH and resuspended in ultra-pure water. DNA was then amplified using primers to detect RasGRP1: 1) PD-SAHR26-Left: 5’-GAGTTCTCTGCTGCCTCCTG-3’; 2) PD-LAHR26-Right: 5’-GGCGGATCACAAGCAATAAT-3’; 3) PD-CAGG-Left: 5’-ATGTCTCCTAGCCGGTGATG. Mutant product was detected at 221 bp, wild type band for the Rosa gene was detected at 341 bp. To detect CRE presence general CRE primers were used with internal control for Globin 1 and Globin 2. CRE: 5’-CACCCTACGTATAGCCG -3’; 5’-GAGTCATCCTTAGCGCCGTA-3’ (product at 300bp).

(S1B) Representative images of papillomas as seen on the ears and tails of *RoLoRiG^+/+^Mx1Cre^+^ mice*.

(S1C) Percentages of GFP^+^ cells in peripheral blood of *RoLoRiG^+/-^ Mx1CRE^+^* mice on days 21 and 94 after pIpC injection. Data are presented as mean ± SEM.

**Supplemental Figure S2: FACS data accompanying Figure 2**

(S2A) FACS schematic to detect T-, B- and myeloid cells.

(S2B-D) Data accompanying Figure 2C. Percentage of T-, B- and myeloid cells in PB assessed by flow cytometry on day 21, 62 and 94 (T cells, n>6; myeloid cells n>8; B cells n>8). pIpC: polyinosinic polycytidylic acid. We observed a small, transient increase in the percentage of total myeloid cells at 21 and 62 days and transient decrease in B cells at 62 days.

(E) T-, B- and myeloid cell percentages in PB of *RoLoRiG^+/-^Mx1CRE^-^* and *RoLoRiG^+/^Mx1CRE^+^* mice at 21 and 94 days after pIpC injection (n>6).

(F) Percentages of GFP^+^ cells within T-, B- and myeloid cell populations in PB of *RoLoRiG^+/-^Mx1CRE^-^* and *RoLoRiG^+/-^Mx1CRE^+^* mice at 21 and 94 days after pIpC injection (n>3).

Data in supplemental Figure S2 are presented as mean ± SEM. Each symbol represents a single mouse. *p<0.05, **p<0.01, ***p<0.001.

**Supplemental Figure S3: *hRASGRP1 overexpression -* bone marrow transfer of LSK cells and analysis of peripheral compartments.**

(S3A) FACS scheme to distinguish recipient (CD45.1/CD45.2 double positive; DP), carrier (CD45.1) and either *RoLoRiG^+/+^Mx1CRE^-^* or *RoLoRiG^+/+^Mx1CRE^+^* donor (CD45.2).

(S3B) Increases in GFP percentage in peripheral blood in recipient mice transplanted with *RoLoRiG^+/+^Mx1CRE^-^* or *RoLoRiG^+/+^Mx1CRE^+^* LSK as a function of time.

(S3C) Mean percentage of recipient (DP), carrier (CD45.1) and either *RoLoRiG^+/+^Mx1CRE^-^* or *RoLoRiG^+/+^Mx1CRE^+^* donor (CD45.2) chimerism in PB of mice at different time points. Congenic-marked (CD45.1) total BM was co-introduced as carrier into CD45.1/CD45.2 lethally irradiated recipient mice.

(S3D) Percentage of T-, myeloid and B-cells in PB of *RoLoRiG^+/+^Mx1CRE^-^* and *RoLoRiG^+/+^Mx1CRE^+^* mice at different time points.

(S3E) Hematopoietic reconstitution was also evident in the spleen with approximately 80% donor contributions in the spleen coming from the CD45.2-marked *RoLoRiG^+/+^Mx1CRE^+^* or *RoLoRiG^+/+^Mx1CRE^-^* LSK cells. Mean percentage of recipient (DP), carrier (CD45.1) and *RoLoRiG^+/+^Mx1CRE^-^* or *RoLoRiG^+/+^Mx1CRE^+^* donor (CD45.2) chimerism in spleens of mice at endpoint.

**Supplemental Figure S4: *hRASGRP1 overexpression -* bone marrow transfer of LSK cells and analysis of bone marrow compartments.**

(S4A) Mean percentage of recipient (double positive-DP), carrier (CD45.1) and *RoLoRiG^+/+^Mx1CRE^-^* or *RoLoRiG^+/+^Mx1CRE^+^* donor (CD45.2) chimerism in BM of mice at endpoint.

(S4B). Bone marrow FACS staining and gating scheme as described, which we combined with congenic CD45 markers and GFP as discriminating tools (**Figure 3**).

(S4C-S4E) Analyses of percentages of LT-HSC, ST-HSC, and MPP cell populations in the BM of recipient mice transplanted with either *RoLoRiG^+/+^Mx1CRE^-^* or *RoLoRiG^+/+^Mx1CRE^+^* LSK (n=9 or more).

(S4F) Percentage of T-, myeloid and B-cells in BM of recipient mice transplanted with either *RoLoRiG^+/+^Mx1CRE^-^* or *RoLoRiG^+/+^Mx1CRE^+^* LSK (n=9 or more).

**Supplemental Figure S5: FACS data accompanying Figure 4**

Examples of three replicates of signal experiments, analyzing p-STAT3 (Y705), p-STAT5 (Y694), p-AKT (S473), p-ERK (T202/Y204), and p-S6 (S235/S236) levels in rested and unstimulated *RoLoRiG^+/+^Mx1CRE^-^* and *RoLoRiG^+/+^Mx1CRE^+^* cells. This panel also depicts the isotype control antibody stainings.

**Cell Sorting of BM-derived CD11B^+^ cells and stimulation assays**

Whole BM cells from either *RoLoRiG^+/+^Mx1CRE^-^* or *RoLoRiG^+/+^Mx1CRE^+^* were harvested at 13-17 weeks after pIpC injection. Single cell suspensions were stained with α-CD11B Ab for 30 mins, on ice and washed twice with FACS buffer. A minimum of 6x10^6^ CD11B+ cells were sorted for cell signaling studies using FACSARIA.

BM-dervied CD11B^+^ sorted cells were rested for 1 hr at 37[Symbol]C in RPMI only. Stimulation of cells was carried out by adding either RPMI with GM-CSF (40ng/ml) or RPMI only as control to the cells for 3 and 10 mins. Following stimulation, cells were fixed in 1.6% PFA at RT for 10 mins. Cells were spun at 500G for 5 mins and permeabilized in ice cold 90% methanol and stored at -20[Symbol]C until staining was performed. For assessment of baseline signaling levels in *RoLoRiG^+/+^Mx1CRE^-^* and *RoLoRiG^+/+^Mx1CRE^+^*, whole BM was fixed in 1.6% PFA and permeabilized in ice cold 90% methanol and stored at -20[Symbol]C until staining was performed.

**Cell signaling assays in baseline BM and phospho-staining**

For Assesment of baseline signaling levels in *RoLoRiG^+/+^Mx1CRE^-^* and *RoLoRiG^+/+^Mx1CRE^+^*, whole BM was harvested as described above and fixed in 1.6% PFA and permeabilized in ice cold 90% methanol and stored at -20. Following permeabilization, cells were rehydrated with FACS buffer and incubated for 5 mins before spinning at 2500g for 5 mins. Cells were then resuspended in FACS buffer and incubated with non-conjugated primary antibodies for pERK, pAKT and pS6, pGSK3 and PE-conjugated antibodies for pSTAT3 and pSTAT5 for 30 mins. Cells were washed and conjugated secondary antibodies were added to the relevant samples for an additional 30 mins. Cells were then washed and at least 10,000 events were acquired in a BD LSRFortessa machine and analyzed using FACSDIVA software. Antibodies information is on Table 1.
